# Hydrogen exchange protection factors can be extracted from sparse HDX-MS data

**DOI:** 10.1101/408559

**Authors:** S. Skinner, G. Radou, R. Tuma, J. J. Houwing-Duistermaat, E. Paci

## Abstract

Hydrogen/deuterium exchange (HDX) monitored by mass spectrometry (MS) is a promising technique for rapidly fingerprinting structural and dynamical properties of proteins. The time dependent change in mass of any fragment of the polypeptide chain depends uniquely on the rate of exchange of its amide hydrogens but determining the latter from the former is generally not possible. Here we show that, if time-resolved measurements are available for a number of overlapping peptides that cover the whole sequence, rate constants for each amide hydrogen exchange (or equivalently, their protection factors) can be predicted. In most cases, the solution may not be unique, so a number of solutions have to be considered. Such analysis always provides meaningful constraints on protection factors and thus can be used in situations where obtaining more refined data is impractical, e.g., high throughput structure and interaction fingerprinting. It also provides a systematic way to improve data collection strategies in order to obtain unambiguous information at single residue level, e.g. for assessing protein structure predictions at atomistic level.

## Introduction

Hydrogen-deuterium exchange probed by mass spectrometry (HDX-MS) (1, 2) is an emerging technique particularly suited to investigate the dynamics of large proteins and their complexes. Dating back to the pioneering work of Linderstrøm-Lang (3), the spontaneous exchange of the amide hydrogens of a protein with deuterium from solvent containing deuterium oxide (^2^H_2_O) has been extensively used to investigate protein folding (4-7). The key to interpreting HDX kinetics is the fact that exchange occurs faster for amides that are solvent-exposed and/or not involved in hydrogen bonds.

Deuterium incorporation has been broadly measured using NMR with residue level resolution (8) for small proteins; for larger proteins and assemblies detection of hydrogen-deuterium exchange by high-resolution mass spectrometry (MS) has emerged as a viable alternative (9, 10). HDX-MS relies on the measurable difference of mass between the deuterated and non-deuterated polypeptide chains. Fragmentation of the protein by proteolysis at low pH and low temperature (i.e., conditions that drastically reduce exchange and thus preserve the isotopic pattern even under non-native conditions, and under which the residual forward and back exchange can be readily corrected for (11, 12)), allows measurement of exchange for specific fragments of the polypeptide chain (usually covering 10-20 amino acids) (13). The deuterium incorporated into side chain groups is rapidly back exchanged and, as a consequence, HDX-MS is only sensitive to the backbone amide exchange.

Monitoring the incorporation of deuterium for each peptide fragment over time yields exchange kinetics; this contains information about local and global stability averaged over all amide NH groups within the peptide. HDX-MS data are usually limited to qualitative analysis, e.g., by mapping the apparent rate of exchange of different peptides on the available structure and comparing the kinetics of the same fragments under different conditions (12, 14, 15).

HDX/MS yields the overall mass change over time for a whole peptide fragment but does not provide direct information on the exchange rate of individual residues. One way to achieve single amide resolution is to obtain fragments that differ by exactly one amino acid and calculating the mass difference. Using enzymatic digestion this is only achievable for a few amino acids under favorable conditions, but recent advances in gas phase fragmentation (2) (e.g., electron capture dissociation (16) and electron transfer dissociation (17, 18)) suggest that HDX-MS can in principle be used to measure hydrogen exchange at single residue resolution. However, even then, gas-phase scrambling (i.e., rapid migration of the incorporated deuterium among backbone and side chains) needs to be minimized by careful optimizing fragmentation conditions which may differ for individual peptide fragments, preventing automated and high-throughput top-down approaches (19). A different strategy, pioneered in Englander’s laboratory, consists in using isotopic envelopes instead of centroid values (20). More recently, it has been shown that combining isotope envelopes with kinetic information a wider time window provides single amide exchange rates for cytochrome c, a 100 amino acid protein (19). However, the required uniform coverage and resolution of isotopic envelopes may be hard to achieve for larger proteins and multi-protein assemblies (18). Only in rare favorable cases, the kinetics and other information have been effectively used to delineate exchange at individual amino acids (21).

In this paper we present a method that extracts individual protection factors from HDX-MS measurements of the mass variation (centroid) of peptic fragments of a polypeptide chain. The method is general, but the degeneracy of the solution (i.e., a set of protection factors for each exchangeable amide hydrogen) depends on the number of peptides (the more the better), on their length (the shorter the better), on the range of times at which the measurement has been measured (the broader the better). The accuracy of the predicted protection factors also depends, obviously, on the accuracy of the measurement itself, although we show here that self-consistent use of the data for overlapping peptides also provides a tool to appraise possible experimental errors in the measurement of the deuterium uptake. We demonstrate that, even in the absence of full, redundant fragment coverage of the protein sequence, and with measurements performed in a relative narrow time window (10 – 10^4^s), the approach presented here provides a relatively small number of solutions, where for a subset of residues the protection factor is uniquely determined, while for others a discreet set of values are possible. We included an option to use isotopic envelopes, if available, to further reduce the number of possible solutions, and uniquely determine the protection factor of each residue also in cases where the information from the centroids in not sufficient for the purpose. These features make the method suitable for analyzing a wealth of existing HDX-MS data and extract crucial information at single-residue level from them and quantifies in a statistically rigorous manner the information contained in the data. When the information in the HDX-MS data provides multiple answers, the tool can be used to guide further experiments that mitigate the underdetermination of the problem.

## Methods

### Principles of hydrogen deuterium exchange probed by mass spectrometry

At neutral pH the exchange is fast for solvent exposed amides while hydrogen bonding, e.g. within helices or β-sheets, slows it down. When fully exposed, the exchange kinetics of the amide is governed by an intrinsic rate, *k*_*int*_ that depends on the temperature, solution pH and side chains of the two neighboring residues (22). Within a folded protein, the exchange of amide hydrogen requires local “opening”of the structure and can be approximated as a two-step process (23):

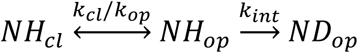

where *k*_*cl*_ and *k*_*op*_ are the local “closing”and “opening”rates. The observed deuterium uptake rate, *k*_*obs*_, can be expressed as:

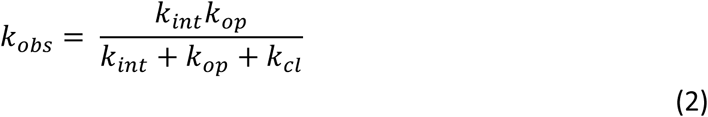

Two limiting regimes, called EX1 and EX2, are invoked in interpreting HDX kinetics of proteins. For both regimes, the protein is considered to be in native conditions, i.e. *k*_*cl*_, ≫ *k*_*op*_. In the EX1 limit *k*_*int*_ ≫ *k*_*cl*,_ implies that the amide exchanges as soon as it becomes exposed to solvent, i.e., *k*_*obs*_ = *k*_*op*_. In this regime the exchange is limited by slow conformational changes that are usually associated with global unfolding (24) or cooperative changes in quaternary structure (15). This regime is readily discerned by a bimodal pattern of isotopic distribution in mass spectra (undeuterated and deuterated species) and by pH independence. In the EX2 limit, *k*_*cl*_≫ *k*_*int*_,

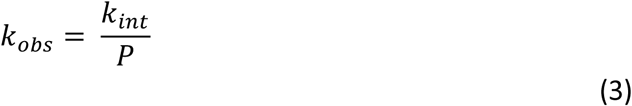

where *p* = *k_cl,_*/*k _op_* is a protection factor for the particular amide hydrogen. The EX2 limit governs exchange under native conditions and is sensitive to local stability. In the EX2 regime the kinetics is sensitive to pH (through *k*_*int*_) and the corresponding isotopic envelope moves progressively to the fully deuterated limit.

HDX-MS measures the change in mass upon deuteration of proteolytic fragments of the polypeptide chain. The deuterium uptake_*DJ*_ for a fragment of polypeptide chain *j*, starting at residue *m*_*j*_ and *n*_*j*_ residues long is

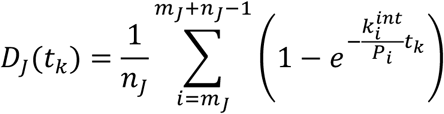

where *p*_*i*_ is the protection factor of residue *i*and *t*_*k*-_ are a set of time points.

### Determination of protection factors from HDX-MS data

The task can be reformulated as determining the set {*p _i_*} so that

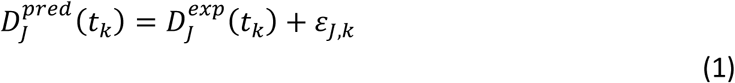

for each *j*and *k*, where *ε*_*j,k*_,-is the error.

This problem corresponds to that of determining the set (or sets) {*p* _*i*_} so that the cost function

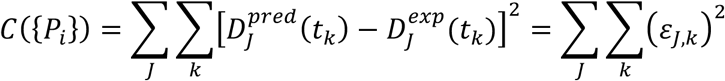

The error originates from the fact that each measured value *D^exp^* is affected by an experimental uncertainty, which is often not reported in the literature. Hence the minimum of the cost function *C*({*p*_*i*_}) is generally not zero. Even in the absence of experimental error, the solution (i.e., sets of protection factors for which *C*({*p*_*i*_}) = 0) is in general not unique. For example, if the deuterium uptake of a single peptide is available, that is long relative to the number of time points at which the experiment has been performed, an infinite number of {*p*_*i*_} sets which minimize *C*({*p*_*i*_}) exist. However, if measurements are available for contiguous overlapping peptides along the polypeptide chain, a unique or a finite number of sets {p _*i*_} for which *C*({*p*_*i*_}) is close to its minimum can be determined for each residue occurring in the contiguous region. The existence and the possible uniqueness of the solution strongly depend on the set of experimental data.

Generating protection factors by drawing from a uniform distribution with boundaries 0 ≤ ln(*p*) ≤ 20 the resulting cost distributed approximately normally with mean 0.62 and standard deviation 0.10. Hence, a random search for solution with cost ~0 is not viable. We opted to perform a random search a number of times (e.g., 10^4^) and use the set with lowest *C*({*P*_*i*_}) as initial condition for a least-square minimization. A sequential least-squares quadratic programming approach, as implemented in SciPy 0.19.1 (25), was used to minimize the cost function.

To favor continuity in the search for minima, we minimized the function

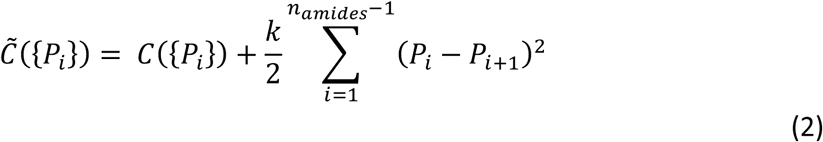

where *k* is a small “elastic constant”and the sum extends to all exchangeable amide hydrogens. For K = 0 the cost is a rugged function with multiple local minima. For larger K (e.g., 10^−4^) the harmonic term added to the cost dominates and prevents this function reaching the lowest possible values of *C*({P_*i*_}). For intermediate values, (e.g., 10^−6^) the harmonic term “facilitates”the identification of sets of solutions that are closest to C = 0. If K = 10^−8^ or smaller the residual harmonic term after minimization is < 10^−5^, i.e., negligible relative to the total residual cost. The lowest minimum (C~0.0051) is reached with K = 10^−8^, and all the solutions (i.e., sets of {P_*i*_}) with C ≤ 0.0055 have been retained as “hits”. The distribution of the values of C sampled in multiple independent simulations with and without harmonic term is shown in Supplementary Information Figure S1.

### Statistical methods

For each residue the median and the interquartile range of the predicted *ln*(P)’s were computed. For various residues with different interquartile ranges histograms were plotted. Since the distributions were multimodal, and the solution for neighboring residues are correlated, we applied model-based clustering method to obtain sets of {*p _i_*} per fragments of the chain. Let *K* be the number of clusters (i.e., the number of solutions for a fragment). Then the following likelihood was used for fragments of length *r*

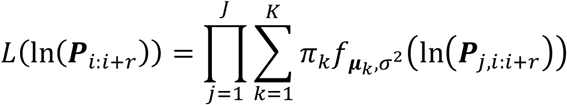

where 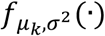 a multivariate normal distribution with mean 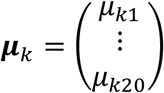 R and variance σ ^*2*^*I* (a *r* × *r* diagonal matrix with σ^*2*^as elements) and π_k-_ the prior probability to belong to cluster *K*. We assume equal variance for the clusters. The log likelihood was maximized by using an expectation maximization (EM) algorithm (26). The number of solutions *K* was chosen based on the lowest Bayesian information criteria (BIC) value. More complex models (different variances) revealed similar results. The choice of the fragment length is somewhat arbitrary and related to the local structure of the protein. Here we chose *r* = 20.

### Calculation of the isotopic envelope

The isotopic-resolved mass spectrum of a peptide (here peptide 75 as an example) is shown in Figure 1; the height of each peak is the natural occurrence of each isotope.

**Figure 1.**
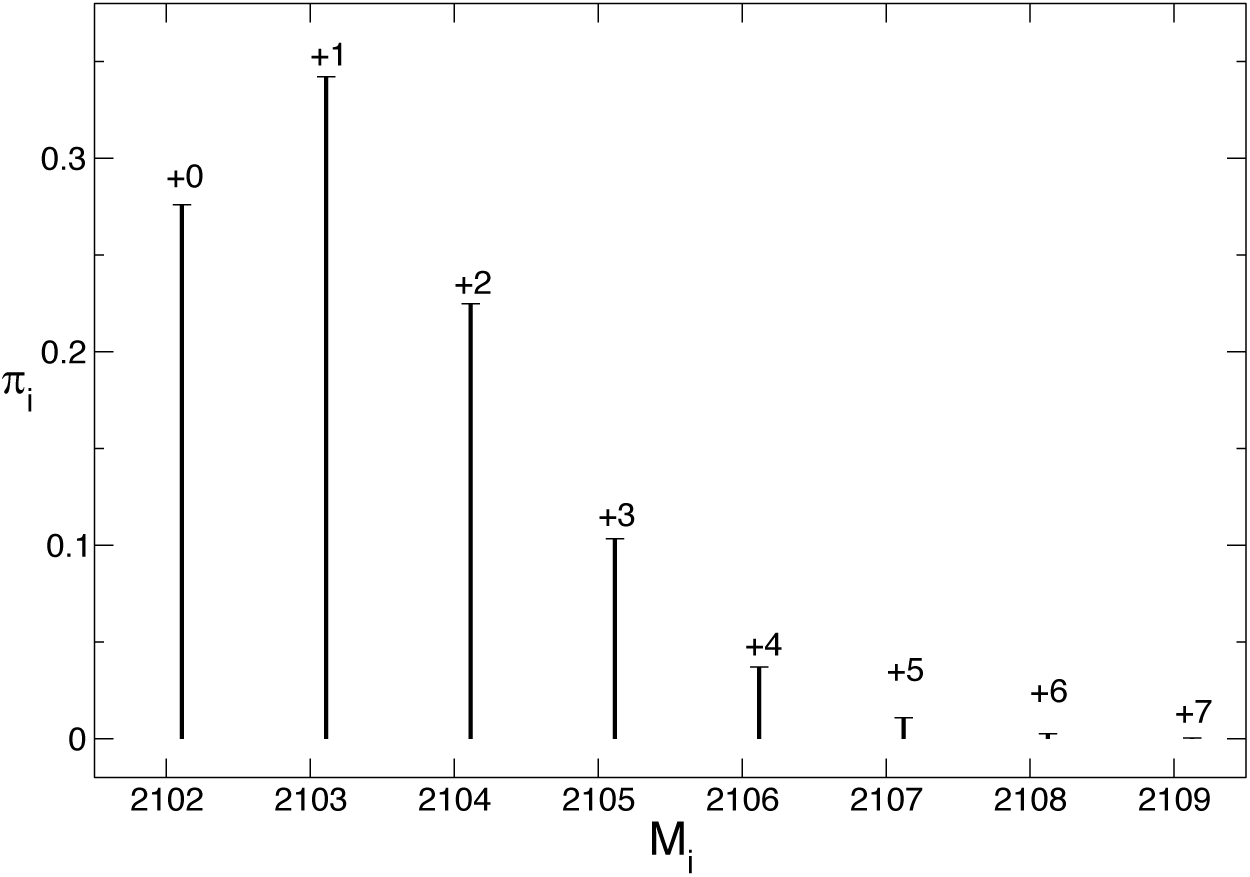
Isotopic envelope for the undeuterated fragment 75. Lines represent the probability associated to the average mass of different isotope numbers.

As amide hydrogen exchange, the intensity of each peak changes with probability depending on the rate of exchange of each amide. For a peptide with *n* exchangeable amides, the probability that *k* (0 ≤ *k*≤ *n*) have exchanged at time *t* is

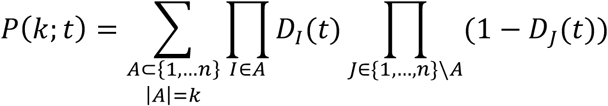

The time dependent isotopic envelopes are calculated simply assigning probability π_*i*_P(K; *t*) to the isotope of mass *m*_*i*_ + (*m*_*D*_ *m*_*H*_)*k* (with *m*_*D*_ –*m*_*H*_ = 1.00627).

The algorithm has been implemented using version 3.6 of the Python programming language using the NumPy, SciPy and Cython libraries. This code is freely available to academics via a GitHub repository (www.github.com). Commercial enterprises can obtain this code subject to a proprietary license.

## Results

We have tested and illustrated this approach on a model protein, C3d, which is a fragment of the complement component C3 (27). C3d is a single-domain protein composed of 297 residues, where residue 1 corresponds to residue 991 of the full C3 molecule. C3d contains twelve α -helices and five 3_10_-helices that are organized into an α-α barrel where most consecutive helices alternate between the inside and the outside of the protein core. Knowledge of the structure, however, is irrelevant and never used in our approach. For C3d HDX-MS measurements (28) for a large number of overlapping polypeptide fragments have been reported (Figure 2). Also important for the reliability of the prediction is that exchange has been measured in a relatively broad time interval (between 10 and 10^4^ s) at seven different time points (19).

**Figure 2.**
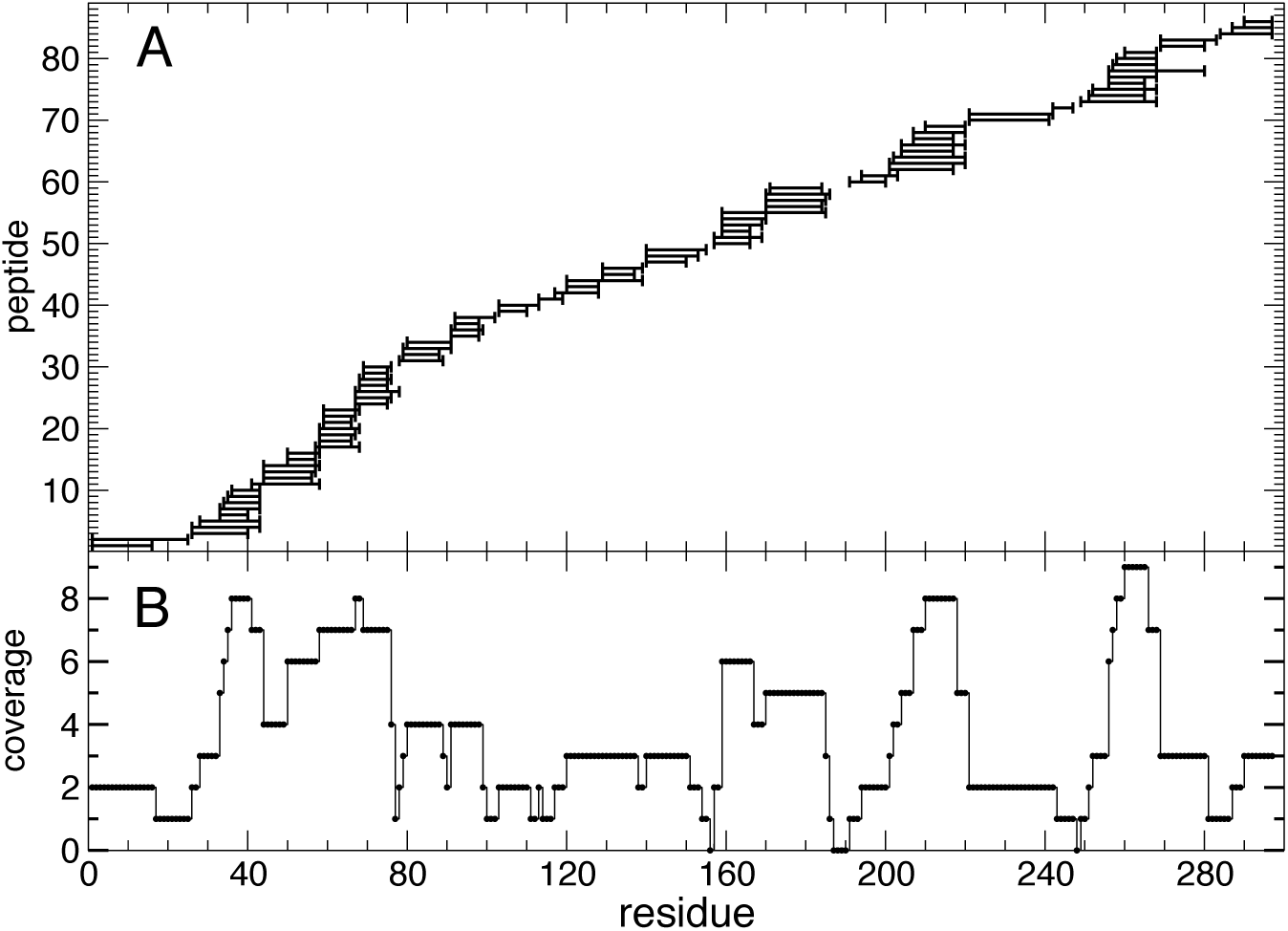
A) Peptides (86 in total) obtained by peptic digestion of the protein C3d studied by HDX-MS by Devaurs *et al.*(28); fragments range between 6 and 26 amino acids in length. B) Coverage, i.e., number of peptides in which an amino acid occurs; of 297 amino acids, only six amino acids are not covered, and their protection factor undetermined. The protein C3d has been chosen as a test case for availability of HDX-MS data with high coverage of the sequence (~98%) and high redundancy (each amino acid occurring in 3.6 independently probed fragments on average).

To determine the values {*p _i_*} that minimize *C*({*p*_*i*_}), we first generate values with a uniform distribution with boundaries 0 ≤ ln(*p*) ≤ 20 (i.e., we assume that the exchange rate of an amide can be as fast as in a completely unstructured peptide and up to 5 × 10^8^ times slower). The process is repeated many times (e.g., 10^4^) and the set with the lowest *C*({*p*_*i*_}) is used as an initial guess for the subsequent minimization (here using the sequential least square quadratic programming method), with constraints 0 ≤ ln(*p*) ≤ 20. The whole procedure is then repeated many times and sets with lowest *C*({*p*_*i*_}) are then selected for further analysis. Details are provided in Methods.

Multiple sets of protection factors {*p _i_*} that represent local minima for *c*({*p*_*i*_}) were found using the procedure described above. Some of these sets show protection factors of adjacent residues being close to opposite boundaries, which are unphysical. Hence, we minimized a slightly modified cost function (Equation 7 in Methods) where a harmonic term is added to disfavor large variations in protection factors between adjacent residues. Including the harmonic term is justified since protection factors correspond to structural properties of the protein (e.g., participation to secondary structure, accessibility to solvent, etc.) which are unlikely to change abruptly between a residue and the previous or next. A small harmonic term prevents reaching local minima and converges to ~0 during the minimization (i.e., the local minimum reached in each minimization is also a minimum of the cost without harmonic term within the tolerance of the minimizer).

Analysis of the sets {*p*_*i*_} that best fit the sets 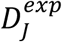is reported in Figures 3 and 4. In Figure 3 the median and interquartile ranges are shown. While for some residues the protection factors are narrowly defined, for others the interquartile ranges show large variations (Figure 3A). Closer examination shows that, in most cases, the distributions of the protection factors for such residues vary narrowly around a discrete set of possible values (see three cases in Figure 3B). The reason that some residues are compatible with different protection factors is clear: there are, in principle, infinite ways in which the same mass variation of a peptide can be obtained by varying the mass of each residue individually, and this increases with the length of the peptide. The uncertainty is restricted by the constraint that the property of each residue is the same in different peptides, but as we have shown (Figure 2) the distribution of fragments is not uniform along the chain as some fragments lack overlap. Consequently, the estimate of a protection factor for a given residue depends on that of neighboring residues. In the case presented here, the whole sequence of the protein is covered redundantly with numerous overlapping peptides. This is the reason why, for several residues, only a single protection factor is compatible with the data and discrete sets of values can be found for the rest.

**Figure 3.**
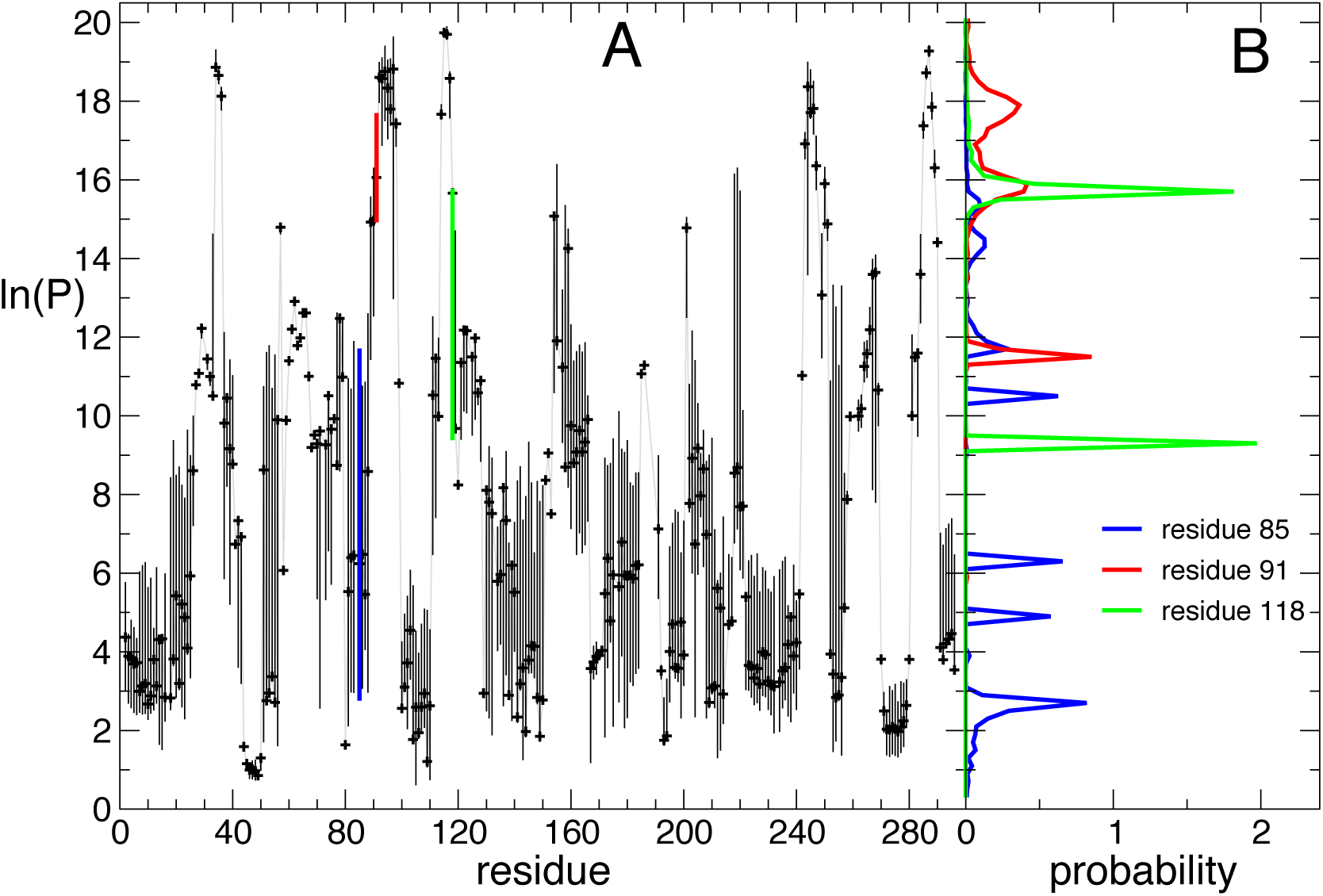
A) Median (black symbols) and interquartile range (black bars) of predicted l*n P*) for all exchangeable amide hydrogen, excluding those not covered by any fragment probed by the HDX-MS measurement (see Figure 2A). Values are computed from over 7,000 independent searches that resulted in a cost *C*<0. 0055. Uncertainty is large for residues poorly covered. An estimation of the distribution of predicted *ln*(*P*) is not possible for all the residues not covered by any fragment (156, 187-190, 248). B) For three positions, the distribution of *ln*(*p*) is shown. B) Histograms of predicted *ln*(*p*) for three exchangeable amide hydrogens with varying uncertainty; the histograms show that the predicted *ln*(*p*) follows a mixture of several Gaussian distributions. The bars representing the interquartile range of the three residues are shown in the same color in A.

**Figure 4.**
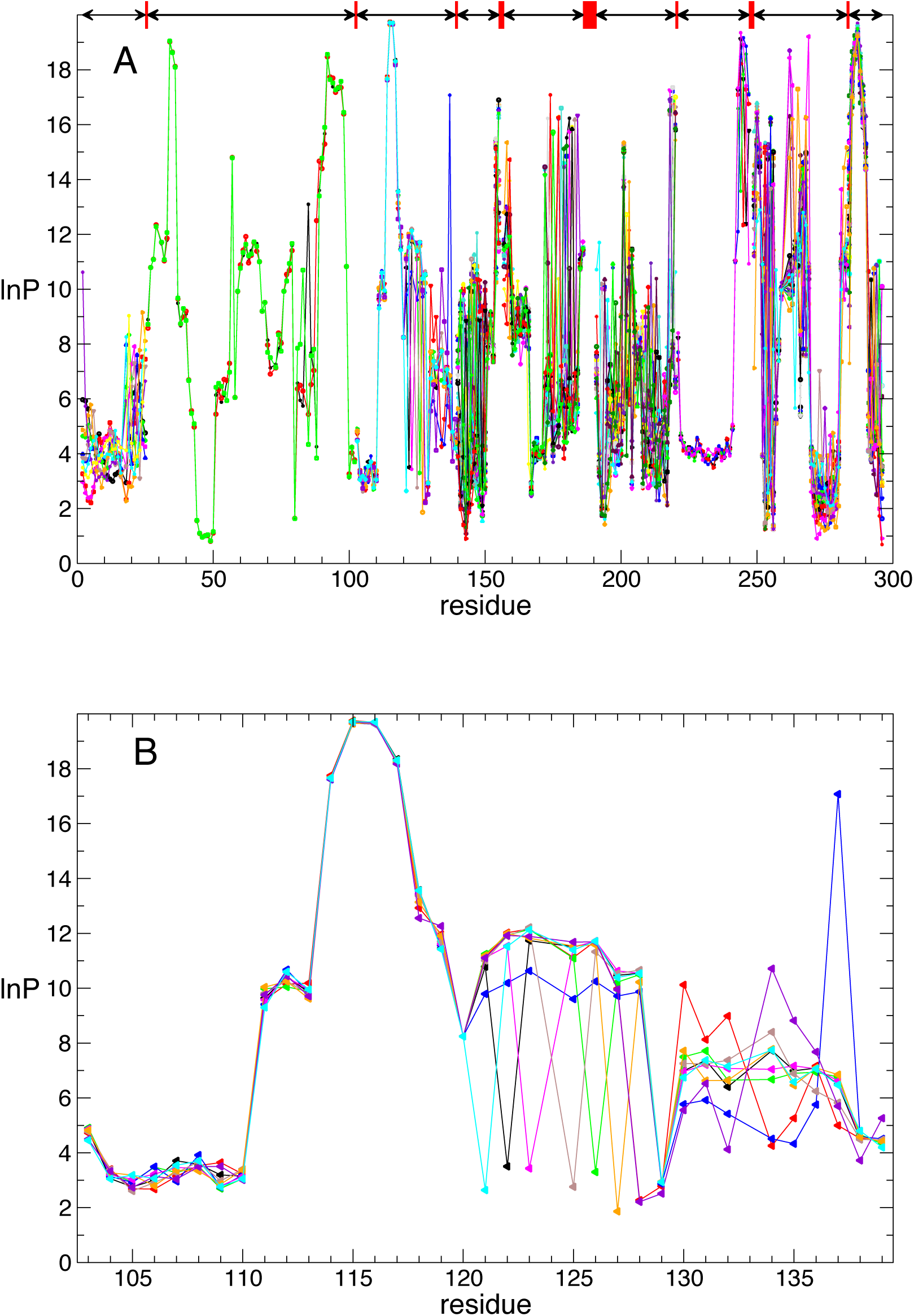
Predicted *ln* (*p*) **from fitting a mixture of multivariate Gaussians. A) For each contiguous region of the polypeptide chain. Contiguous regions, i.e., regions of the polypeptide chain covered by overlapping peptides, are represented with arrows; gaps between contiguous regions are represented by red boxes. The multivariate means of each cluster of solutions are shown, in different colors. B) The nine different clusters for the contiguous region encompassing residues 103-139.**

The interdependence of the prediction of the protection factor for a residue with that of neighboring residues can be assessed by fitting mixtures of multivariate Gaussians. Interdependence extends as far as contiguous stretches of the polypeptide chain covered by overlapping peptides. In the case presented here, there are nine contiguous regions, that have been fitted independently. The multivariate means for all contiguous region of the polypeptide chain is depicted in Figure 4A. The multivariate means for a single contiguous region (residue 103-139) is shown if Figure 4B. For some residues the protection factor is uniquely defined, while for others alternative values are possible, but only a limited number of their combinations. Such a representation of the statistical analysis of the solutions also helps to discriminate regions with well constrained protection factors from those with considerable ambiguity and also gives upper and lower bounds on possible values.

### Comparison with experimental deuterium uptake data

As presented above, multiple sets of protection factors can satisfy the experimental data equally well. The experimentally measured and predicted deuterium uptake of six different peptides is shown in Figure 5. Different sets of protection factors fit equally well the experimental data but deviate from each other outside the measured time window. For each peptide, the 10 sets of protection factors with lowest cost (among the ~7000 sets with cost *C*< 0.0055) have been used to calculate the deuterium uptake over a time interval covering 10 orders of magnitude. For peptides 1, 71 and 82 we observe that experimental data is perfectly reproduced in the time interval probed by the experiment, but different sets of predicted protection factors provide alternative profiles for times shorter than those measured experimentally. This is related to the fact that the three peptides fall in a region of low coverage (see Figure 2), but also highlights the necessity of measurements at shorter times (< 1*s*). The opposite is true for peptide 37, where the experimental data only cover times at which the exchange has not yet occurred, and the variability in the predicted protection factors is responsible for a different time dependence of the predicted deuterium uptake at long time (> 10^6^s). The prediction of the deuterium uptake for peptide 20 appears to be robust at all timescales, most likely due to the information contained in the considerable number of peptides that partially overlap with it. The deuterium uptake of peptide 6 represents the most eminent case in which the fit is consistently suboptimal, and possible reasons discussed below.

**Figure 5.**
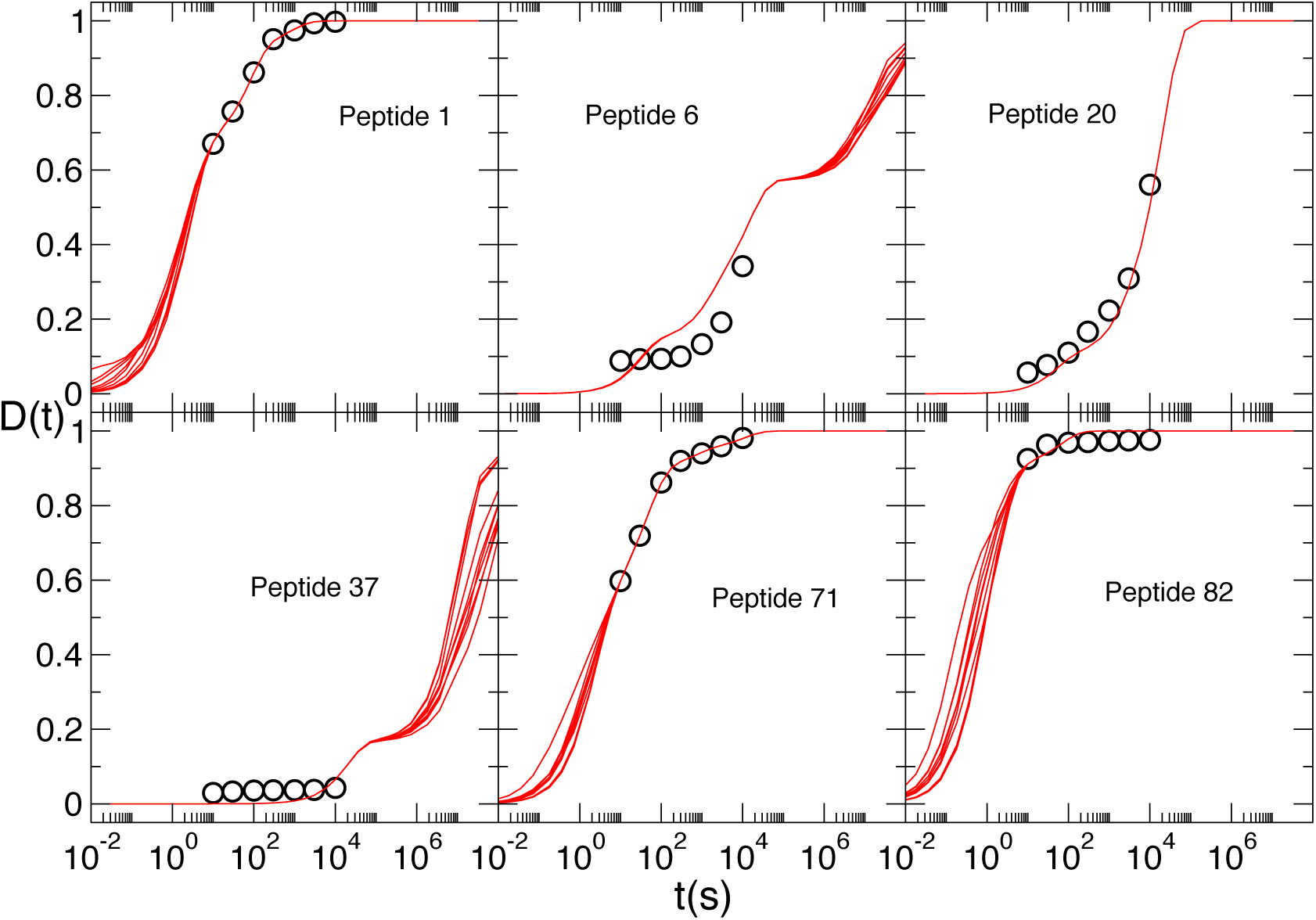
**D**^exp^ (*t*_*k*_) **(circles) and *D***^pred^ (*t*) **(continuous lines) as a function of time for six of the 86 peptides experimentally probed. Continuous lines are three fits among the thousands with a total cost *C***<0.0055.

The root mean square deviation (averaged over all the different sets of predicted protection factors) between experimental 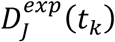 and calculated D_J_ (t_k_)is shown in Figure 6. Also shown is the deviation between experiment and model when the cost function is minimized for individual peptides (in which case, the protection factors of individual residues are severely underdetermined; an example is provided in SI Appendix, Figure S5): a value larger than zero highlights that the experimental deuterium uptake cannot be fitted exactly with the sum of exponentials in Equation (3); this provides a lower limit of the experimental error in the measurement. Most peptides can be fitted exactly individually, but not when considered simultaneously (this is for example the case of the highly overlapping peptides 56-59 that all include residues 171-183) because of experimental errors on the deuterium uptake of a peptide and/or that of partially overlapping of flanking peptides.

**Figure 6.**
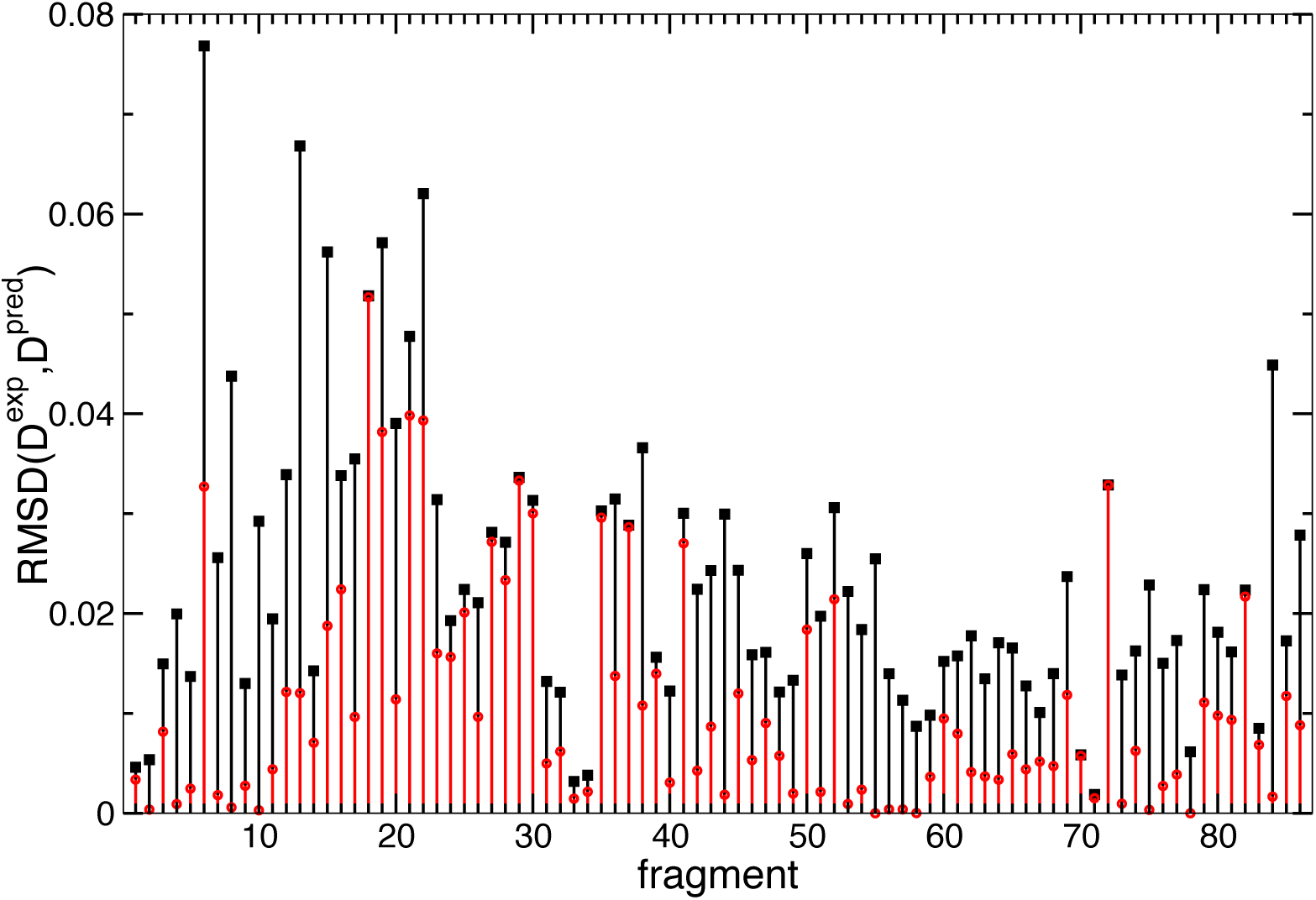
Deviation between experiment and calculation for each peptide. In black is shown the deviation between model and experimental data when the deuterium uptake of all peptides is optimized simultaneously. In red the same deviation when each peptide is fitted individually; a value different from zero means that no set {*p* _*i*_} can be found that perfectly fit the experimental deuterium uptake, corresponding to a lower limit for the average experimental error for that peptide. Deviation is largest for peptide 6 also highlighted in Figure 5. For most other peptides the deviation is small and within the presumed statistical error of the experimental measurement.

### Prediction of the isotopic envelope

The knowledge of protection factors (or equivalently of the time dependent deuterium uptake of individual amides) allows the prediction of the time evolution of the isotopic envelope of each peptide. For a peptide that shows large variation in the prediction of the protection factor for a subset of amino acids (Figure 7A), the isotopic envelope is shown at different times in Figure 7B, where different colors refer to different patterns of protection factors, all of which reproduce exactly the experimental deuterium uptake for the whole peptide. Thus, when isotopic envelopes are available a quantitative comparison can help to further disambiguate the alternative sets of protection factors.

**Figure 7.**
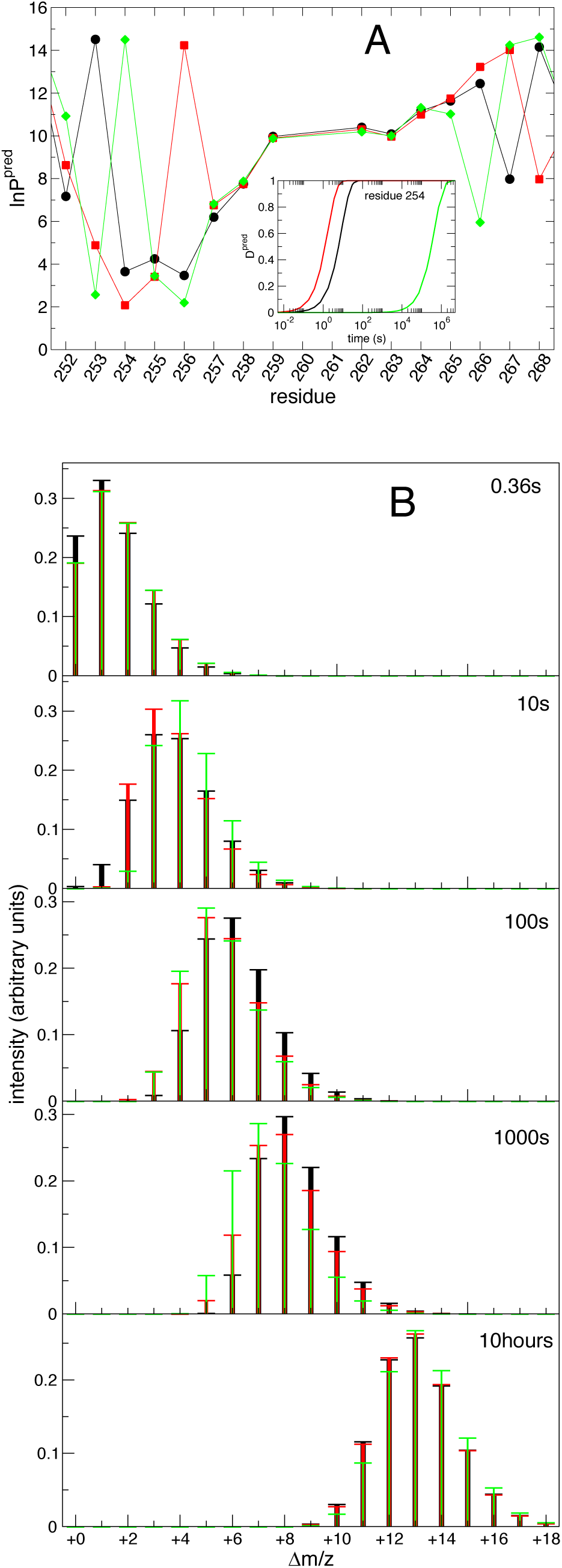
A) Alternative patterns of protection factors for all non-proline residues of peptide 75 encompassing residues 252-268; in the inset are shown the alternative deuterium uptake profiles for residue 254 for the alternative sets of protection factors. B) Isotopic envelope of peptide 75 at five different time points; the monoisotopic species has ***m***/*z* =2102.11.

## Discussion

A large number of studies using HDX-MS report results for a set of peptides resulting from the enzymatic digestion of proteins. Results are most often reported as deuterium uptake curves (measured as centroid mass) or “butterfly charts”and thus provide only qualitative information at the resolution level of several amino acids. Limited attempts to extract rates as single residue level have been previously presented (29, 30).

In this paper we have presented a method that can estimate HDX protection factors for individual residues from HDX-MS centroid mass measurements as a function of time. Our approach provides alternative sets of protection factors that agree with the experiment. For several residues the prediction provides a set of unique protection factor values. For others, we obtain a range of solutions that, in most cases, is well represented by a relatively small set of discrete values. These considerations depend strongly on the set of experimental data. The existence of a unique or of finite sets of solutions depends on the length and overlap of the peptide fragments. Also, peptides that exchange much faster than ~10s or slower than 10^4^s, which is the interval usually probed, provide little or no information.

The contribution of statistical and systematic errors to the experimental data deserves further discussion. Statistical error could be easily included in the present approach, by accepting all the solutions that satisfy the experimental data within the error. Systematic errors depend mostly on the correction for back (and sometimes also forward) exchange. A phenomenological correction for back exchange could be included in the present approach by minimizing the function 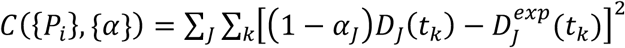where *α*_j_ is the fraction of deuterated amides that exchanges back to hydrogen, i.e., assuming that the ratio of exchange and back exchange events is constant over all time points. Another simple extension of the method consists of adding a weight to the fit of individual fragment, in the case in which the experimental uncertainty is known for each fragment. To demonstrate the working of our method, we used a dataset where fragments redundantly cover most of the protein sequence. The result is not a single set of protection factors, but a family of them. This is inevitable when fragments are long and coverage of the sequence uneven (in Supplementary Information Figures S3 and S4 we show how the results would change if the case in which fragments were shorter, denser, or both).

In contrast to previous methods (29, 30), we chose to generate a large number of solutions and perform a thorough statistical analysis of them. The median and interquartile range provide information on residues for which a safe prediction of the protection factors can be made, and those for which it cannot. For those residues where the prediction is ambiguous, we show that, in most cases, the distribution of the prediction is multimodal, with values distributed narrowly around discrete set of possible values. A multivariate analysis provides the best guess of the protection factors for adjacent residues, since the prediction for one residue depends on the prediction for the others that appear in the same peptide, or in peptides that continuously overlap. The method proposed here represents a search for self-consistent solutions to the problem and provides statistical analysis of the outcomes, thus yielding the best guess and associated uncertainty of the protection factors that reproduce the time course of the mass change measured experimentally. The approach primarily uses centroid data which represents the vast majority of the published HDX-MS results, hence it is widely applicable. Availability of protection factors for each residue can be converted structural information (31-34) and used to generate or validate models of proteins and complexes with unknown structure.

The method proposed here also gives an indication of how the uncertainty could be reduced. For example, it highlights regions of the polypeptide chain where a different fragmentation approach would provide additional constraints and reduce the uncertainty. Where the time evolution of the isotopic envelope is available for specific peptides, it provides a practical way to deconvolute the spectra and reduce or eliminate the uncertainty on the estimation of the protection factors of the peptide and reduce that of contiguous peptides. Given the increased availability of mass spectrometers and automation of the exchange and fragmentation process, the method presented here has strong potential to turn a mostly qualitative analysis into a quantitative one.

## Author Contributions

S.S., R.T., E.P., G.R. and J.J. H.-D. designed and performed the research, S.S., E.P. and J.J. H.-D. analyzed the data, S.S., R.T., E.P. and J.J. H.-D. wrote the article.

## Acknowledgments

We thank Didier Devaurs for sharing with us the complete datasets in Ref(28). We acknowledge financial support from Wellcome Trust grant number 096686/Z/11/Z.

